# Altered stimulus-specific adaptation at the auditory cortex of a mouse model of Fragile X Syndrome

**DOI:** 10.64898/2026.09.09.750387

**Authors:** Abdullah Abdullah, Xiuping Liu, Jun Yan, Ning Cheng

**Affiliations:** Cumming School of Medicine, University of Calgary, Calgary, T2N 1N4, Canada; Faculty of Veterinary Medicine, University of Calgary, Calgary, T2N 1N4, Canada; Hotchkiss Brain Institute, University of Calgary, Calgary, T2N 1N4, Canada; Alberta Children’s Hospital Research Institute, University of Calgary, Calgary, T2N 1N4, Canada; Owerko Centre, University of Calgary, Calgary, T2N 1N4, Canada; Department of Physiology and Pharmacology, University of Calgary, Calgary, T2N 1N4, Canada

**Keywords:** Auditory hypersensitivity, adaptation, Fragile X Syndrome, auditory cortex, electrophysiology study

## Abstract

Auditory hypersensitivity or decreased sound tolerance is a common phenotype of Fragile X Syndrome (FXS). Impairments in adaptation, defined as the reduction in the neuronal responsiveness to repeating sounds, can contribute to this prevalent phenotype. Previous studies on event-related potential observed impairments in mismatch negativity (MMN) in both FXS individuals and in the *FMR1*-knockout (KO) mouse model of FXS. Therefore, in the present study we characterized stimulus-specific adaptation (SSA), a neural correlate of MMN, at the auditory cortex (AC) of anesthetized (with ketamine/xylazine) female and male postnatal day 20 (P20) wild-type (WT) and *FMR1*-KO mice, using the oddball paradigm with either 4 Hz or 1 Hz repetition rate. We observed robust SSA at the 4 Hz repetition rate of the oddball paradigm in the AC neurons of all four groups of animals. We also noted that the strength of SSA diminished between 4 to 1 Hz repetition rate. In addition, at both the 4 Hz and 1 Hz repetition rate, reduced SSA was observed particularly in the male *FMR1*-KO mice compared to their WT counterparts, while female WT and *FMR1*-KO mice displayed similar SSA. In terms of sex differences, male *FMR1*-KO mice had lower SSA than female *FMR1*-KO mice, while SSA was similar in male and female WT mice. Overall, our observation of reduced SSA in the male *FMR1*-KO mice suggests that impairments in neuronal adaptation potentially contribute to the auditory hypersensitivity phenotype particularly within the male population.

## Introduction

Auditory hypersensitivity is a consistent phenotype of various neurodevelopmental disorders including autism spectrum disorder (ASD) (Ida-Eto et al., 2017). Fragile X Syndrome (FXS) is a common monogenic cause of ASD (Niu et al., 2017), which occurs due to the silencing of the Fragile X Messenger Ribonucleoprotein 1 (*FMR1)* gene and subsequent absence of the Fragile X Messenger Ribonucleoprotein (FMRP) (Liu et al., 2018). Consequently, this leads to various FXS symptoms such as anxiety, social deficits, language impairments and decreased tolerance to sound (Abdullah et al., 2025; Möhrle et al., 2026; Niu et al., 2017). Although auditory hypersensitivity is a prevalent phenotype, very little is known about its underlying mechanisms hindering the development of effective therapeutic interventions.

Repetitive sounds like raindrops are common in the natural environment. The auditory system is responsible for filtering out irrelevant or repetitive sounds from the surrounding environment while responding to unusual or novel sounds, a phenomenon called adaptation (Lanting et al., 2013). Adaptation is commonly involved with processes such as change detection (Pérez-González & Malmierca, 2014), wherein an increased responsiveness is observed to rare tones compared to repeated tones. In humans, change detection can be observed in cortical electroencephalography (EEG) studies as mismatch negativity (MMN), using the oddball paradigm where a stronger negative deflection of the auditory event-related potential (ERP) is evoked by the novel sound compared to the frequent sound (Nieto-Diego & Malmierca, 2016). In FXS individuals, MMN was observed to be decreased in males between 18 and 42 years of age (Van Der Molen et al., 2012), whereas another study with male and female FXS patients between ages 7 and 34 years old observed enhanced MMN amplitudes (Proteau-Lemieux et al., 2025). Another study by Ethridge et al observed no difference in MMN amplitude in male and female FXS patients aged 4-51 years old compared to the control group (Ethridge et al., 2020).

Although the findings were inconsistent, previous studies indicated that neural adaptation could be impaired in FXS. MMN is generally considered a scalp-recorded potential reflecting the summed activity of several cortical areas (Nelken & Ulanovsky, 2007), and therefore cannot pinpoint the specific neural generators of adaptation. The closest phenomenon to MMN on a neuronal level is stimulus-specific adaptation (SSA). SSA is a stimulus-specific response, wherein neuronal response decreases following a series of repeated sounds but remains highly responsive to a rare sound (Wang et al., 2014). Although SSA is a well-defined phenomenon in the primary auditory cortex (AC) (Ulanovsky et al., 2003), to our knowledge previous studies did not characterize SSA in the *FMR1*-knockout (KO) mouse model, which is a well-established mouse model of FXS and syndromic autism (Ahmed et al., 2026; Wang et al., 2026; Zhan et al., 2020).

In the present study we investigated whether impaired SSA at the AC during early auditory development could contribute to auditory hypersensitivity in FXS. Robust SSA was observed in both the P20 wild-type (WT) and *FMR1*-KO mice at the 4 Hz repetition rate. Interestingly, we observed that SSA diminished with increasing interstimulus intervals in all the groups. Finally, at the 4 Hz repetition rate, we observed significant genotype difference in SSA only in the male *FMR1*-KO mice compared to their WT counterparts.

## Methodology

### Animals

Wild-type (FVB.129P2-Pde6b+ Tyrc-ch/AntJ, Jax stock No: 004828) and *FMR1-*KO (FVB.129P2-*Pde6b*^+^ *Tyr^c-ch^ Fmr1^tm1Cgr^*/J, Jax stock No: 004624) mice were obtained from Jackson Laboratory (ME, USA) and maintained at the mouse housing facility in Health Sciences Animal Resources Centre, University of Calgary. Mice were bred at the animal facility of the University of Calgary. Pups were weaned at P20 and group-housed (five mice per cage) with same sex littermates. Mouse cages were housed at the mouse facility with automated 12-hr light and dark cycle and provided with food (standard mouse chow) and water as necessary. Electrophysiology recordings were conducted between 09:00h and 19:00h. All procedures were carried out within the guidelines outlined by the Canadian Council for Animal Care and were also approved by the Health Sciences Animal Care Committee at the University of Calgary.

### Surgery

The mice were first provided with an intraperitoneal injection of ketamine (85 mg/kg) and xylazine (15 mg/kg) to anesthetize the mice throughout surgery and electrophysiological recording. Further maintenance doses of ketamine/xylazine (17 and 3 mg/kg, respectively) were provided throughout this period, based on responsiveness to tail pinching. The mouse’s head was mounted onto a custom-made head holder inside a sound-proof chamber with electromagnetic proofing, and the mouse’s head was further clamped between the palate and nasal bones. The scalp, subcutaneous tissue and muscle were then carefully removed to expose the skull. The position of the mouse’s head was further adjusted to align the bregma and lambda of the skull with the horizontal plane. A dental drill was used to perform a small craniotomy measuring ∼3mm in diameter to make an opening at the left auditory cortex (2.2–3.6 mm posterior to bregma, 4–4.5 mm lateral to the midline) based on coordinates from the Allen Brain Atlas.

### Recording at the AC

A ∼2-MΩ impedance tungsten wire microelectrode was lowered perpendicularly with a hydraulic microdrive into the left auditory cortex. Signals from the microelectrode were passed to a RA16PA multichannel preamplifier and a RA16 Medusa multichannel amplifier (Tucker-Davis Technologies). Spike signals were recorded through one channel amplified 10,000 times, filtered by a bandpass of 0.3-10 kHz. Another channel recorded local field potential (LFP) amplified 1000 times and filtered with a bandpass of 1 to 200 Hz. Subsequently, the signals were digitized at a sampling rate of 25 kHz and stored using the BrainWare data acquisition software (Tucker-Davis Technologies, Inc., Gainesville, FL, USA). Tone-evoked responses were observed 400 to 700 μm below the brain surface.

### Acoustic stimulation

Pure tones were digitally generated and converted to analog sinusoidal waves by an RZ6 MULTI I/O processor (Tucker-Davis Technologies, Inc., Gainesville, FL, USA). The analog signals were transferred via a digital attenuator and presented through a speaker (MF1, Tucker-Davis Technologies., Gainesville, FL, USA) positioned 45° and 15 cm from the right ear of the mouse. The tone sound level (expressed as dB SPL) of the loudspeaker was calibrated at the same position, prior to the experiment, using a condenser microphone (Model 2520, Larson-Davis Laboratories, USA) and a microphone preamplifier (Model 2200C, Larson-Davis Laboratories, USA). BrainWare data acquisition software was used manually or digitally to deliver tones.

### Sampling receptive fields of AC neurons

Once stable tone-evoked responses were observed, a frequency-amplitude scan (FA-scan) was performed to record the excitatory responses of the AC neurons to pure tones (20 ms, 5 ms rise and fall time) of different frequency and sound level. The FA-scan consisted of a series of tones delivered at frequencies that were equally spaced on a logarithmic scale, and at sound levels that ranged from 10 to 90 dB SPL in 10 dB increments. The inter-stimulus interval between each tone was 250 ms, and the entire session lasted for approximately 10 minutes. The receptive fields or frequency tuning curve for each AC unit were obtained from the FA-scan data.

### Data processing of FA-scan

All spike data recorded by the BrainWare acquisition software was saved as DAM files. These files were then processed by our custom-made data processing software, SoundCode. Next, the trigger level was set manually at 20% greater than average background amplitude to detect tone-evoked neuronal firing or spikes. Spike number was defined as the sum of spikes to five identical stimuli within a 70 ms window, that is 5 ms post tone onset to 75 ms post tone onset. Minimum threshold (MT) was determined from the FA-scan of the AC neurons and was defined as the lowest sound intensity to elicit responses of the AC neurons across frequencies.

### Oddball paradigm

The oddball paradigm consisted of three conditions with varying numbers of standard and deviant tones. Two tones f_1_ and f_2_ were selected from the FA-scan within 10-30 dB above MT. In the first condition an equal number of f_1_ and f_2_ tones (60 ms) were provided to ensure that neuronal firing was similar to both tones. The second condition (Condition 2) delivered f_1_ with 90% probability (as standard tones) and f_2_ with 10% probability (as deviant tones) in the stimulus sequence. In the third condition (Condition 3), the probability of f_1_ and f_2_ in the stimulus sequence was reversed. This set of conditions ensured that the AC neurons could correctly identify deviant tones solely due to the stimulus probability, and not due to the confounding features of the stimulus (such as different frequencies). The two frequencies were provided at a frequency contrast of Δf= 0.37 (corresponding to frequency ratio of 0.526) where Δf = (f_2_-f_1_)/ (f_2_*f_1_)^1/2^, and repetition rates of both 2 and 4 Hz (Malmierca et al., 2009). In addition, each tone in the stimulus sequence was repeated 40 times and provided randomly within each condition **(Figure 1A)**.

**Figure 1:**
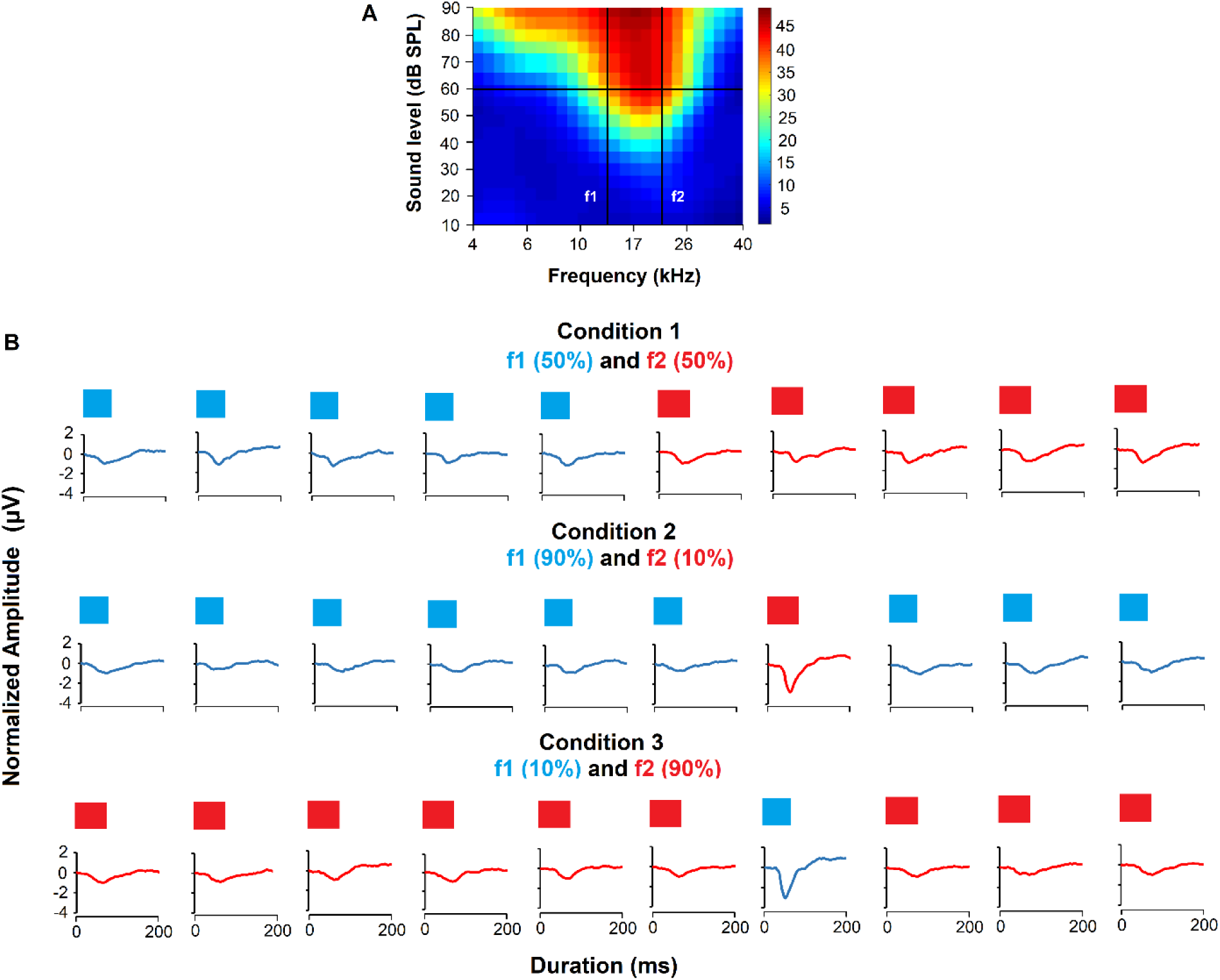
(A) Representative contour plot of a frequency-amplitude scan (FA-scan) from a single AC neuron showing two tones (f_1_ and f_2_) that were selected for the oddball paradigm. Two vertical lines indicate the two frequencies, and the horizontal lines indicate that the sound intensity was selected 30 dB above the minimum threshold. Color scale represents spike counts. (B) Oddball paradigm methodology: In Condition 1 of the oddball paradigm, f_1_ and f_2_ had an equal probability of occurring in the sequence. Condition 2 delivered f_1_ with 90% probability and f_2_ with 10% probability within the stimulus sequence. The probability of f_1_ and f_2_ was reversed in Condition 3. In this figure and all the following figures, high probability and low probability tones will be referred to as standard and deviant tones respectively. Representative LFP traces are shown for all conditions.

### Data and statistical analysis

The amplitude of the negative component of the LFP was measured as neuronal response to the tones in the oddball paradigm. For the SSA analysis, ST(f_1_) and ST(f_2_) denote the neuronal responses to f_1_ and f_2_ when each frequency was provided as the standard tone, while DT(f_1_) and DT(f_2_) denote the neuronal responses to f_1_ and f_2_ when each frequency was provided as the deviant tone. Accordingly, ST(f_1_) and DT(f_1_) were obtained from Condition 2 and 3, respectively, whereas DT(f_2_) and ST(f_2_) were obtained from Condition 2 and 3, respectively. Next, two indices were calculated namely common SSA index (CSI) and frequency specific index (SI).

1. Common SSA index (CSI): The CSI was calculated as a frequency-independent measure of the overall difference between responses to deviant and standard tones, in both Condition 2 and 3. The following equation was used: CSI= [DT(f_1_) + DT(f_2_) - ST(f_1_) - ST(f_2_)] / [DT(f_1_) + DT(f_2_) + ST(f_1_) + ST(f_2_)].
2. Frequency-specific index (SI): The frequency-specific index (SI) was analyzed to quantify SSA separately for each frequency by comparing the neuronal response to the same frequency when it was presented as a deviant tone versus when it was presented as a standard tone. The following equation was used: SI = [DT (f_i_) - ST (f_i_)] / [DT (f_i_) + ST (f_i_)] to quantify deviant and standard tone responses to f_1_ and f_2_.

The values of both indices range between −1 and +1, with +1 indicating increased responsiveness to deviant tones and −1 indicating increased responsiveness to standard tones (Malmierca et al., 2009). Statistical tests were performed in GraphPad Prism 10.2.3 (GraphPad Software, San Diego, California), and figures were generated in GraphPad Prism and MATLAB (MATLAB R2023b, MathWorks, Inc.). All statistical analyses are presented in the figure legends. All data are shown as Mean ± SEM, except the scatterplots, and α was set at 0.05.

## Results

### Robust SSA was observed with 4 Hz repetition rate at the AC of both the WT and the *FMR1*-KO mice

A total of 41 and 42 AC neurons were analyzed in the wild-type (WT) (Male= 20, Female= 21) and *FMR1*-KO (Male= 23, Female= 19) mice, respectively, at the 4 Hz repetition rate. We first analyzed whether stimulus-specific adaptation (SSA) can be observed at the AC of both the WT and *FMR1*-KO mice. We quantified frequency-specific index (SI) of the AC neurons to f_1_ and f_2_ separately and presented the data in the form of scatterplots **(Figure 2)**. Majority of the AC neurons in both the male and female WT and *FMR1*-KO mice occupied the upper right quadrant, with the mean located on the 45° dotted line, indicating robust SSA to both frequencies in the oddball paradigm **(Figure 2 A-D)**. In addition, by analyzing the distribution of common SSA index (CSI) **(Figure 2 E-H)** we observed that the CSI of most AC neurons was skewed towards the positive value suggesting that SSA can be reliably observed in all the groups.

**Figure 2:**
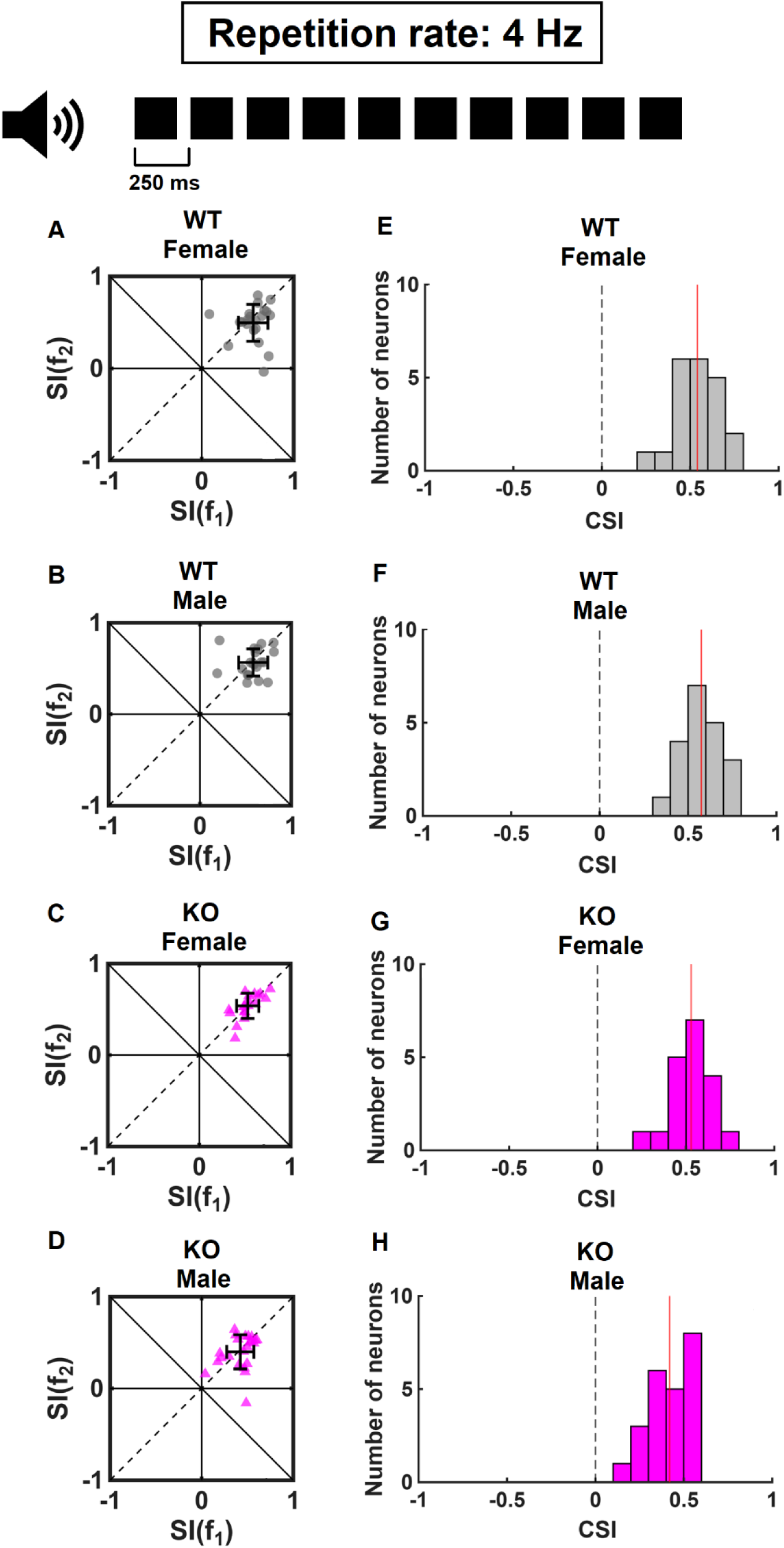
SSA at the 4 Hz repetition rate of the oddball paradigm. (A-D) Scatter plot showing SI, a frequency-specific measure of SSA, to f_1_ vs f_2_. In all scatter plots, top-right quadrant indicates the area within which AC neurons show strong SSA to both frequencies. Bottom-left quadrant indicates the area within which neurons show no SSA. Top-left and bottom-right indicates areas within which AC neurons show SSA to only a specific frequency. The black crosses show the mean and standard deviation (SD) for both axis. (E-H) Distribution of CSI in all the groups. Red line indicates the mean of the distribution.

### SSA weakened with increase in interstimulus interval in both the WT and the *FMR1*-KO mice

At the 1 Hz repetition rate, we analyzed a total of 37 WT neurons (Male= 19, Female= 18) and 35 *FMR1*-KO neurons (Male= 19, Female= 16). In terms of SI, most of the AC neurons in both the male and female WT and *FMR1*-KO mice occupied the upper right quadrant and were also clustered near the center **(Figure 3 A-D)**. Further analysis of the CSI distribution **(Figure 3 E-H)** showed that most of the AC neurons in all the groups were located close to zero, suggesting that SSA may be diminished at the 1 Hz repetition rate.

**Figure 3:**
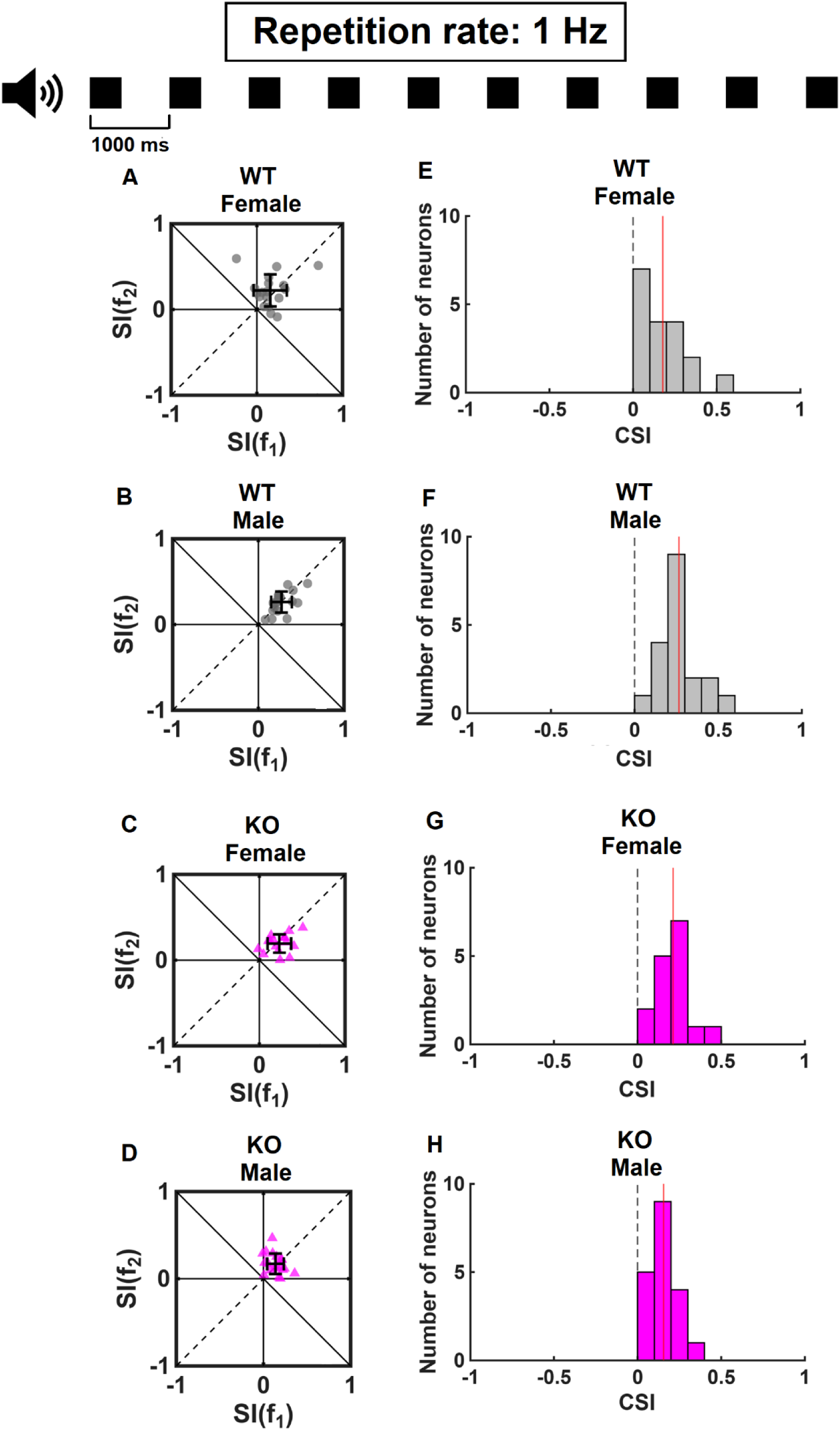
SSA at the 1 Hz repetition rate of the oddball paradigm. (A-D) Scatter plot showing SI to f_1_ vs f_2_. In all scatter plots, top-right quadrant indicates the area within which AC neurons show SSA to both frequencies. Bottom-left quadrant indicates the area within which neurons show no SSA. Top-left and bottom-right indicates areas within which AC neurons show SSA to only a specific frequency. The black crosses show the mean and standard deviation (SD) for both axis. (E-H) Distribution of CSI in all the groups. Red line indicates the mean of the distribution.

Therefore, we also analyzed whether SSA changed with repetition rate in all the groups. Repetition rate was observed to significantly influence the strength of SSA in all the groups. SSA was lower at 1 Hz compared to 4 Hz repetition rate in the WT male (p= <0.0001), WT female (p= <0.0001), *FMR1*-KO male (p= <0.0001) and *FMR1*-KO female (p= <0.0001) groups **(Figure 4 A-D)**.

**Figure 4:**
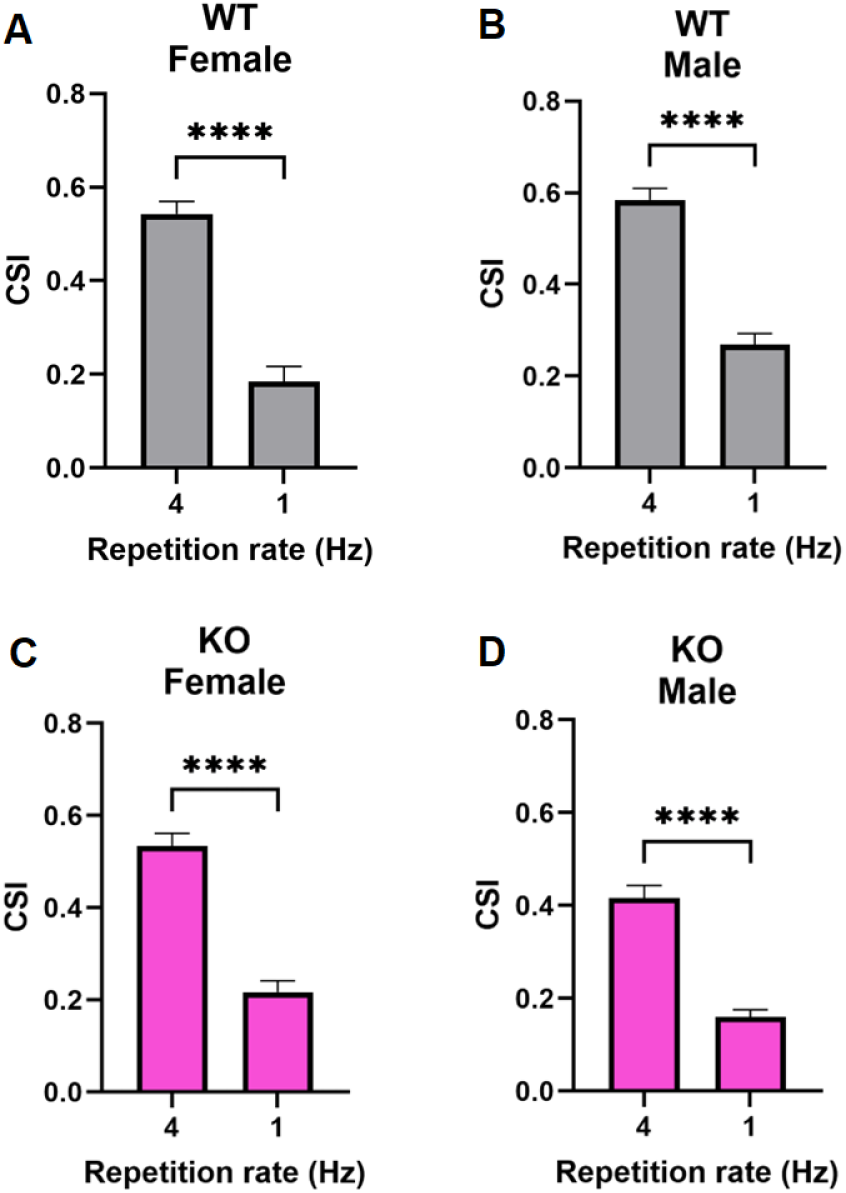
CSI between different repetition rates in the oddball paradigm. Comparison of CSI between 4 and 1 Hz repetition rate in the female and male WT mice (A&B), as well as in the female and male *FMR1*-KO mice (C&D). Analysis: Non-parametric unpaired Mann-Whitney test (****: p <0.0001).

### Decreased SSA was observed at the AC of male P20 *FMR1*-KO mice

Finally, we compared the strength of SSA at the AC of WT and *FMR1*-KO mice. At the 4 Hz repetition rate, a significant genotype difference was observed in CSI that took into account of both Condition 2 and 3 (p= 0.0013), with further post-hoc analysis indicating that CSI was significantly decreased in the male *FMR1*-KO mice compared to their WT counterparts (p= 0.0001). Although the sex effect was non-significant (p= 0.1547), significantly reduced CSI was also observed in the male *FMR1*-KO mice compared to the female *FMR1*-KO mice (p= 0.0122) **(Figure 5C)**. At the 1 Hz repetition rate, no main effects of genotype was observed in CSI (p= 0.1342), with further post-hoc analysis indicating that CSI was significantly decreased in the male *FMR1*-KO mice compared to their WT counterparts (p= 0.0122) **(Figure 6C)**.

**Figure 5:**
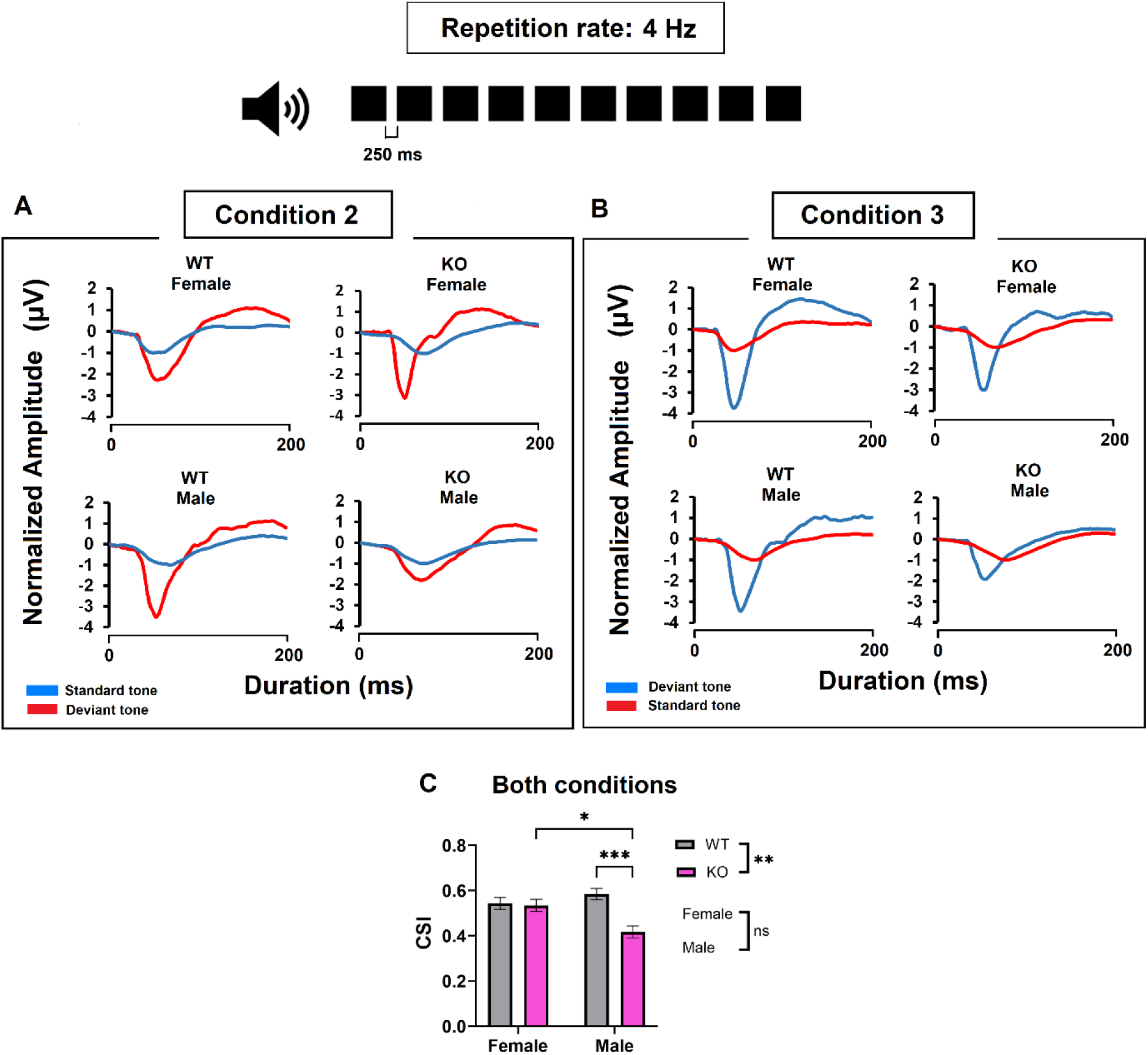
SSA magnitude in both the female and male WT and *FMR1*-KO mice, at the 4 Hz repetition rate. (A) Representative LFP traces from both female and male WT and *FMR1*-KO mice in Condition 2. (B) Representative LFP traces from both female and male WT and *FMR1*-KO mice in Condition 3. (C) CSI comparison between genotype in the male and female WT and *FMR1*-KO mice within both oddball conditions. Analysis: Ordinary Two-way ANOVA with post-hoc Tukey’s analysis (*: p <0.05, **: p <0.01, ***: p <0.001).

**Figure 6:**
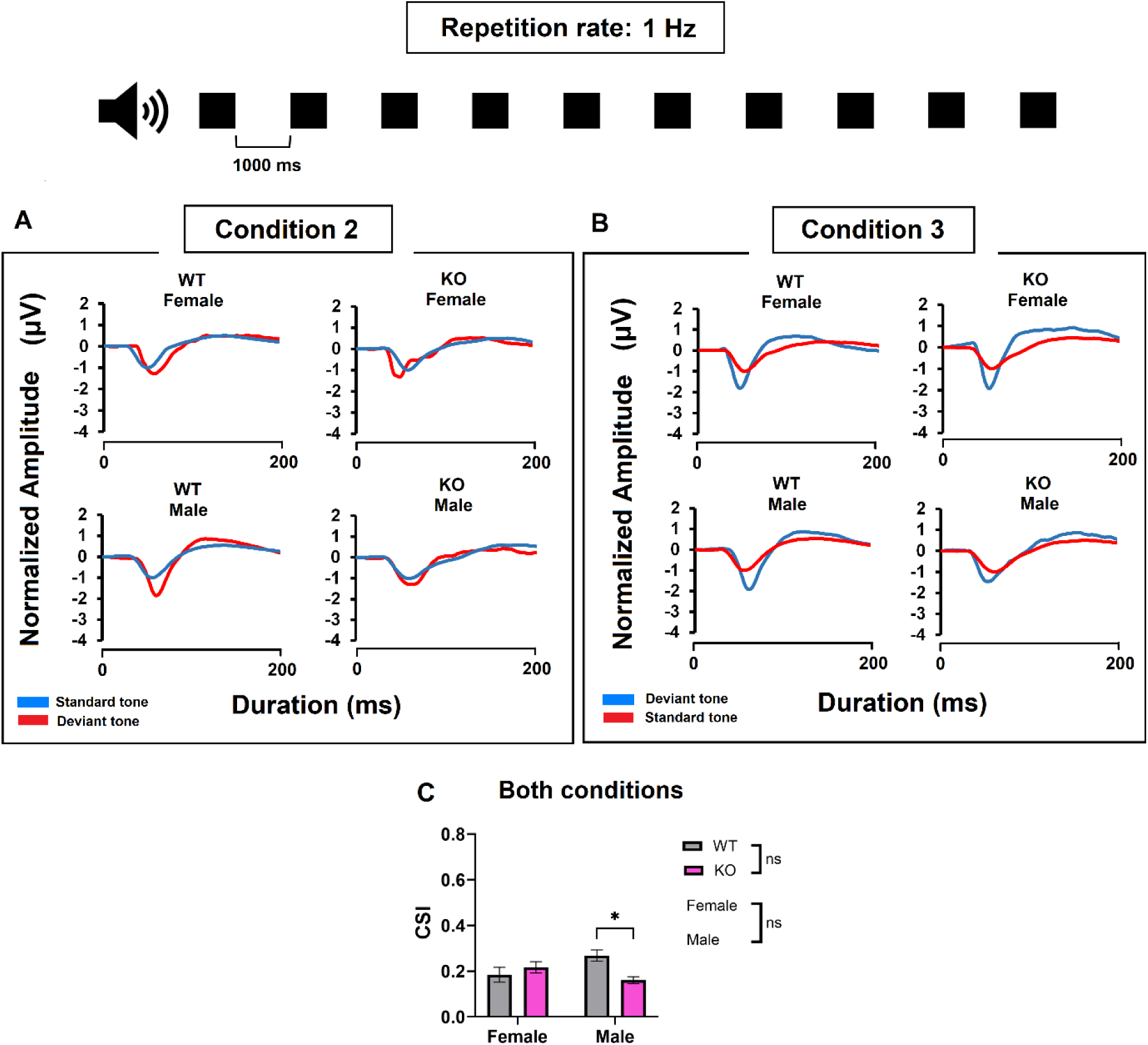
SSA magnitude between genotypes in both the female and male WT and *FMR1*-KO mice, at the 1 Hz repetition rate. (A) Representative LFP figures from both female and male WT and *FMR1*-KO mice during Condition 2. (B) Representative LFP figures from both female and male WT and *FMR1*-KO mice during Condition 3. (C) CSI comparison between genotype in the male and female WT and *FMR1*-KO mice within both oddball conditions. Analysis: Ordinary Two-way ANOVA with post-hoc Tukey’s analysis (*: p <0.05).

## Discussion

In the present study, we performed *in-vivo* electrophysiology recordings at the auditory cortex (AC) of postnatal day 20 (P20) wild-type (WT) and *FMR1-*knockout (KO) mice. AC neurons in both the WT and *FMR1-*KO mice displayed robust stimulus-specific adaptation (SSA) at the 4 Hz repetition rate **(Figure 2)**. At both the 4 Hz and 1 Hz repetition rate, diminished SSA was observed in the male *FMR1*-KO mice compared to their WT counterparts **(Figure 5 and 6)**. At the 4 Hz repetition rate, the male *FMR1*-KO mice exhibited lower SSA compared to the female *FMR1*-KO mice **(Figure 5)**. Overall, our study observed robust SSA in all mice groups along with significant genotype and sex differences in SSA at the AC.

### Strength of SSA was dependent on the repetition rate

The SSA magnitude was observed to be greater at 4 Hz compared to the 1 Hz repetition rate in all the groups **(Figure 4)**. This was consistent with a previous study which investigated membrane potential response to standard and deviant tones at the rodent AC, and observed that SSA diminished with increasing inter-stimulus intervals (ISI) (between 300 and 1200 ms) (Hershenhoren et al., 2014). Synaptic depression is responsible for suppressing neuronal responses to repeating tones with ISI beyond 100 ms (Wehr & Zador, 2005). Therefore, with slower repetition rates or longer ISI, AC neuronal responses to standard tones may recover more following synaptic depression, resulting in decreased SSA at the 1 Hz repetition rate.

### SSA is significantly reduced in male *FMR1*-KO mice

A major finding of this study was that SSA was significantly reduced in male *FMR1*-KO mice compared to male WT mice at both the 4 Hz and 1 Hz repetition rates **(Figures 5 & 6)**. This impairment is likely related to dysfunction of GABAergic inhibitory interneurons at the AC, which are known to play a critical role in shaping SSA. Within the GABAergic population, two principal subtypes namely the parvalbumin-expressing (PV+) and somatostatin-expressing (SOM+) interneurons comprise approximately 40% and 18% of cortical GABAergic neurons respectively (Xu et al., 2010) and contribute to SSA through complementary but distinct mechanisms (Natan et al., 2015).

Of relevance to the male-specific SSA impairment, SOM+ interneurons selectively suppress excitatory responses to frequently occurring (standard) tones, without affecting responses to rare (deviant) tones, thereby directly amplifying the difference between standard and deviant responses (Natan et al., 2015). A reduction in SOM+ interneuron activity at the AC of male *FMR1*-KO mice could therefore directly account for the reduced SSA observed in the present study. While direct evidence for SOM+ deficits specifically at the AC of *FMR1*-KO mice is currently lacking, this remains a plausible mechanism. It should be noted that Kalinowska et al found that cell type-specific deletion of *FMR1* from SOM+ neurons did not cause behavioral deficits or affect global protein synthesis in the cortex or hippocampus (Kalinowska et al., 2022), suggesting that FMRP loss from SOM+ neurons alone may not drive broad FXS-like pathology. Further studies are required to directly assess whether SOM+ expression or function is selectively impaired at the AC of male *FMR1*-KO mice.

PV+ interneurons are also an important component of the inhibitory network shaping auditory cortical processing. Wen et al. reported that PV+ neuron density was reduced specifically in the male *FMR1*-KO mice at the AC at P21 (Wen et al., 2018), driven by elevated MMP-9 activity that impairs perineuronal net (PNN) formation around PV+ cells (Dziembowska et al., 2013; Ethell & Ethell, 2007; Lensjø et al., 2017). Similarly, Kalinowska et al. also observed that *FMR1* deletion at PV+ neurons resulted in behavioral deficits along with enhanced de novo protein synthesis in the hippocampus of adult (2-6 months old) *FMR1-*KO mice (Kalinowska et al., 2022).

PV+ interneurons provide broad, non-specific inhibitory gain control at the AC (Moore & Wehr, 2013), and their loss may contribute to the E-I imbalance and auditory hypersensitivity characteristic of *FMR1*-KO mice. Regarding SSA specifically, Natan et al. originally reported that PV+ suppression increased responses to standard and deviant tones equally, predicting no net effect on SSA (Natan et al., 2015). However, a more recent study by Yarden et al. found that PV+ suppression facilitated deviant responses approximately ten times more than standard responses (Yarden et al., 2022), suggesting that PV+ dysfunction could also contribute to altered SSA under certain conditions. The exact contribution of PV+ deficits to the SSA impairment in male *FMR1*-KO mice therefore warrants further investigation.

### SSA is not significantly different between WT and female *FMR1*-KO mice

In contrast to males, SSA was not significantly different between female WT and *FMR1*-KO mice at either repetition rate **(Figure 5 & 6)**, suggesting that the inhibitory circuit mechanisms underlying SSA are relatively preserved in females. PV+ interneurons are known to exhibit sex-dependent differences in density, excitability, and connectivity across cortical regions (Seney & Joffe, 2026). Therefore, the mechanisms regulating PV+ maturation including perineuronal net formation and MMP-9 activity may be differentially regulated in female mice, potentially conferring resilience against the effects of FMRP loss.

On a similar note, the preserved function of the PV+ interneurons in females is consistent with preserved SSA but does not by itself fully explain the preserved SSA. Whether SOM+ interneurons are also spared in female *FMR1*-KO mice, which could account for intact SSA, remains an open question. The mechanistic basis for the sex difference in SSA in *FMR1*-KO mice is therefore not fully resolved, and future studies directly comparing inhibitory interneuron subtypes between male and female *FMR1*-KO mice at the AC are needed.

### Conclusion

Auditory hypersensitivity is a common and prevalent phenotype of FXS. Previous EEG/ERP studies observed alterations in mismatch negativity (MMN) in FXS (Proteau-Lemieux et al., 2025; Van Der Molen et al., 2012); however whether adaptation impairments at the neuronal level could also be attributed to auditory hypersensitivity was unknown. Our study demonstrated that the degree of SSA, a neural correlate of MMN, at the AC of male *FMR1*-KO mice was lower compared to that of the male WT mice during early auditory development. SSA was preserved at the AC of female *FMR1*-KO mice during this period, which is potentially an important methodological consideration for future studies. Future studies can investigate SSA in adult mice to observe whether impaired SSA can be observed later in auditory development in the male and female *FMR1*-KO mice.

## Author Contribution

AA participated in designing the experiment, data analysis, interpretation, writing and revising the manuscript. XL participated in designing the experiment, recording and data analysis, and editing and revising the manuscript. JY participated in designing the experiment, data analysis and interpretation, and editing and revising the manuscript. NC participated in designing the study, data analysis and interpretation, and editing and revising the manuscript.

## Abbreviations

FXS: Fragile X Syndrome
ASD: Autism spectrum disorder
AC: Auditory cortex
*FMR1*: Fragile X Messenger Ribonucleoprotein 1
FMRP: Fragile X Messenger Ribonucleoprotein
*FMR1*-KO: *FMR1*-knockout
WT: Wild-type

