## Supplementary Table for "Altered stimulus-specific adaptation at the auditory cortex of a mouse model of Fragile X Syndrome"

### Supplementary materials

**Table S1:** Statistical analysis for **Figure 5C** to compare genotype and sex in the oddball paradigm at the 4 Hz repetition rate. Analysis: Ordinary Two-way ANOVA followed by post-hoc Tukey's test (*: p <0.05, **: p <0.01, ****: p <0.0001, ns: non-significant). Green indicates significant p-value.

<table>
<tr><th>Source of Variation</th><th>P value</th><th>P value summary</th><th>F (DFn, DFd)</th></tr>
<tr><td>Sex</td><td>0.1547</td><td>ns</td><td>F (1, 79) = 2.065</td></tr>
<tr><td>Genotype</td><td>0.0013</td><td>**</td><td>F (1, 79) = 11.06</td></tr>
<tr><td colspan="4">Post-hoc Tukey's Analysis</td></tr>
<tr><td colspan="2">Group comparisons</td><td colspan="2">P-value</td></tr>
<tr><td colspan="2">Female WT vs Female FMR1-KO</td><td colspan="2">0.9957</td></tr>
<tr><td colspan="2">Male WT vs Male FMR1-KO</td><td colspan="2">0.0001</td></tr>
<tr><td colspan="2">Female WT vs Male WT</td><td colspan="2">0.6916</td></tr>
<tr><td colspan="2">Female KO vs Male KO</td><td colspan="2">0.0122</td></tr>
</table>

**Table S2:** Statistical analysis for **Figure 6C** to compare genotype and sex in the oddball paradigm at the 1 Hz repetition rate. Analysis: Ordinary Two-way ANOVA followed by post-hoc Tukey's test (*: p <0.05, **: p <0.01, ****: p <0.0001, ns: non-significant). Green indicates significant p-value.

<table>
<tr><th>Source of Variation</th><th>P value</th><th>P value summary</th><th>F (DFn, DFd)</th></tr>
<tr><td>Sex</td><td>0.5823</td><td>ns</td><td>F (1, 68) = 0.3054</td></tr>
<tr><td>Genotype</td><td>0.1342</td><td>ns</td><td>F (1, 68) = 2.298</td></tr>
<tr><td colspan="4">Post-hoc Tukey's Analysis</td></tr>
<tr><td colspan="2">Group comparisons</td><td colspan="2">P-value</td></tr>
<tr><td colspan="2">Female WT vs Female FMR1-KO</td><td colspan="2">0.8042</td></tr>
<tr><td colspan="2">Male WT vs Male FMR1-KO</td><td colspan="2">0.0122</td></tr>
<tr><td colspan="2">Female WT vs Male WT</td><td colspan="2">0.0815</td></tr>
<tr><td colspan="2">Female KO vs Male KO</td><td colspan="2">0.3948</td></tr>
</table>
